# A Novel Simulation Framework for Validation of Ecological Network Inference

**DOI:** 10.1101/2023.08.05.552122

**Authors:** Erik Kusch, Anna C. Vinton

## Abstract

1. Understanding how the differential magnitude and sign of ecological interactions vary across space is vital to assessing ecosystem resilience to biodiversity loss and predict community assemblies. This necessity for ecological network knowledge and their labour-intensive sampling requirements has spurred the creation of ecological network inference methodology. Recent research has identified inconsistencies in networks inferred using different approaches thus necessitating quantification of inference performance to facilitate choice of network inference approach.
2. Here we develop a data simulation method to generate data products fit for network inference and subsequently quantify the validity of two well-established ecological interaction network inference methods – HMSC and COOCCUR. The simulation framework we present here can be parameterised using real-world information (e.g., biological interactions observed in-situ and bioclimatic niche preferences) thus representing network inference capabilities in real-world applications. Using this framework, it is thus possible to evaluate the performance of any ecological network inference approach.
3. We identify a concerningly large range in accuracy of inferred networks as compared to true, realisable association networks. These differences in inference accuracy are governed by a paradigm of input data types and environmental parameter estimation as previously suggested. To establish a workflow for quantification of network inference reliability, we suggest analysis procedures with which to explore inference and detection probabilities of association types of different identity and sign with respect to bioclimatic niche preferences and association strength of association partner-species.
4. With this study, we provide the groundwork with which to validate and compare ecological network inference methods, and ultimately vastly increase our ability to understand and predict species biodiversity across space and time.

## INTRODUCTION

Despite their complexity (e.g., in directionality, sign, and strength), biological interactions (e.g., competition, mutualism, predation) can be described readily – even among and across large assemblages of interacting species – via ecological networks. Ecological networks quantify the direction and sign of interactions (i.e., links) between species (i.e., nodes) as either positive, negative, or absent while also describing their strength/magnitude (Morales-Castilla et al., 2015). Real-world counterparts of such abstracted biological interactions and their respective networks are undergoing changes at local and macroecological scales simultaneously, but traditional in-situ quantifications become intractable across large scales due to sampling effort requirements (Jordano, 2016). Nevertheless, it is at these macroecological scales that climate change impacts ought to be evaluated (Lavorel & Garnier, 2002; Rapacciuolo & Blois, 2019; Wang et al., 2016) and so macroecological networks have to be inferred from proxies. This has led to the development of interaction inference approaches, most of which utilize spatial patterns of species co-occurrence (e.g.; Griffith, Veech, & Marsh, 2016; Blonder & Morueta-Holme, 2017; Damgaard, Ehlers, Ransijn, Schmidt, & Svenning, 2018; Lamonica, Pagel, & Schurr, 2021; Swain et al., 2021; Tikhonov et al., 2021; Weiss-Lehman et al., 2021; Bimler, Mayfield, Martyn, & Stouffer, 2022) to identify ecological networks. Due to their demonstrated usefulness in studying and predicting (1) community assembly patterns (Freilich et al., 2018; Ovaskainen et al., 2017), (2) ecosystem processes (Dongli et al., 2022) and services (Keyes et al., 2021) as well as (3) ecosystem stability (Korkmazhan & Dunn, 2022; Thébault & Fontaine, 2010), it is pivotal that networks inferred via these approaches are accurate representations of realised biological interactions.

A recent comparison of network inference outcomes has demonstrated that – even when analysing the same ecosystem – inferred networks are inconsistent across inference approaches and geographic scales (Kusch et al., 2023). Thus, practitioners of ecological network inference need to carefully consider which network inference approach to use as inferred networks differ considerably across scales and approaches. To streamline choice of inference methodology for specific study purposes, criteria of inference approach applicability have been proposed (Kusch et al., 2023). However, objective measurements of inference performance to judge quality of ecological network inference remain absent. To account for the ongoing development of network inference approaches, a generalizable inference validation framework that can take into account ongoing developments is vital.

To study performance of ecological network inference approaches, we require a framework which produces testable networks and data with which to carry out inference of networks which can then be compared to the underlying testable networks. While one previous attempt has been made at creating such a comparison framework (Lavender et al., 2019), this comparison arguably suffers from a lack of biological realism which has also been identified as a set of pitfalls in a recent overview of ecological network inference practices (Blanchet et al., 2020). First, limiting data simulations and network inference to undirected links (i.e., biological associations) fails to represent a large part of the ecological interaction type spectrum (Harris, 2016). To study the effect of link directionality on ecological network inference, a network inference performance assessment framework must therefore be capable of generating synthetic networks containing directed (i.e., interactions) or undirected (i.e., associations) links. Second, neglecting to simulate and infer data according to bioclimatic niche preferences may lead to overestimations of the impact of ecological interactions (Turnbull et al., 2013). Thus, a simulation framework for network inference validation must explicitly consider bioclimatic preferences of ecological network-contained species to represent more realistic biodiversity products for network inference. Third, spatial biodiversity products for network inference beyond co-occurrence records (i.e., abundance or performance matrices) have been demonstrated to lead to vastly different inference outcomes even when using the same underlying statistical framework for network inference (Kusch et al., 2023). Conclusively, to quantify the accuracy-changes in network inference owing to differences in spatial input types, the simulation framework must be capable of generating biodiversity products beyond co-occurrence. Finally, to mirror real-world data as closely as possible, such a simulation framework ought to be capable of incorporating considerations of population dynamics such as density dependence and dispersal as these factors have been demonstrated to shape spatial patterns of biodiversity alongside biological interactions (Saupe et al., 2015).

Here we present a novel simulation-based toolbox for ecological network inference performance assessments overcoming the above-mentioned limitations. This framework centres on a spatially explicit, individual-based simulation for the generation of data products ready for network inference. With this framework, we address the impact of bioclimatic niches and biological interactions as well as density dependence and dispersal dynamics in shaping species distributions and abundance through time and space. This data generation framework can be parameterised either with real-world network and species data (e.g., species-specific bioclimatic niches and dispersal potential), or randomly generated network and species data (for which we provide supporting functionality). While the former will allow for precise quantification of network inference performance given specific study systems, the latter enables a more general assessment of network inference reliability across a range of systems and species. Because of the flexibility of the data input and robustness of the mathematical simulation approach, this framework is generalisable across study ecosystems and species pools. Moreover, this simulation framework provides a flexible toolbox with which additional considerations beyond the ones presented here may be addressed (e.g., different types of species interactions, environmental change impacts). Using this simulation framework, we subsequently carry out ecological network inference using two well-established approaches of ecological network inference – HMSC (Ovaskainen et al., 2017) and COOCCUR (Veech, 2013). While the former allows us to assess the performance of network inference while accounting for bioclimatic preferences and spatial products beyond simple co-occurrence records, the latter inference approach is blind to climatological gradients and restricted in input data to co-occurrence matrices. We expect that network inference will more accurately identify true pairwise associations when bioclimatic preferences are accounted for and data products corresponding to co-abundance or co-performance rather than co-occurrence are utilised, thus making HMSC inference more accurate. Finally, we propose a generalisable framework with which to study the likelihood of inference and detection (i.e., correct inference) of different types on pairwise links. Ultimtely, the framework for data generation and process of network inference assessments we present here represent an advancement in our understanding of performance and applicability of ecological interaction inference methodology.

## MATERIAL & METHODS

### ECOLOGICAL NETWORK MATRICES

When encoding ecological networks, it is common practice to represent nodes (i.e., species) and their links (i.e., interactions or associations) as matrices (Ávila-Thieme et al., 2023; Opedal & Hegland, 2020). Within these matrices, columns and rows represent species-identities while cells contain the link weight (interaction/association strength) between corresponding species pairs. Network inference approaches can detect network matrices as either representing directed (Bimler et al., 2022) or undirected networks (Morueta-Holme et al., 2016; Tikhonov et al., 2020; Veech, 2013).

Directedness of ecological network links is identified through arrangement of values within the corresponding network matrix. Undirected, association matrix entries are contained either in the upper or lower triangle of the species-by-species matrix (i.e., every species in an association pair affects the partner equally). Directed, interaction matrices, on the other hand, contain the interaction strength and sign of an actor species (stored in columns) onto a subject species (stored in rows) and vice versa. While undirected network matrices identify biological associations (e.g., probability of co-occurrence of two species), directed network matrices allow for the identification of more complex biological processes like predator-prey interactions. Conclusively, for our framework of network inference validation presented here, we have implemented support for interaction and association matrices.

While our simulation framework can be parameterised with ecological network matrices as observed in nature, we provide functionality for creation of random network interaction or association matrices. Optional arguments of this helper function control maximum value of link strengths and sparsity of the matrix (i.e., how many of all possible values should be exactly 0) representing an absence of interaction or association between species.

### SIMULATION FRAMEWORK

In order to generate data to assess network inference performance, we developed a spatially explicit, individual-based birth-death model. Here, we iteratively simulate birth and death events of individuals belonging to interacting species (as defined via a network matrix) across a two-dimensional (*x* and *y*), continuous landscape.

### THE MODEL

Individuals in this model have constant birth rates (*b*_0_), while their death rate over time (*d*(*t*)) is determined by their (1) population density and carrying capacity (P), (2) environmental maladaptation (*E*), and (3) interaction effects of neighbouring individuals (Ω).

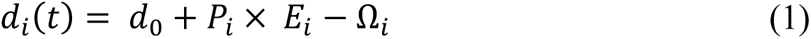

Where *d*_*i*_(*t*) is the dynamic death rate for individual *i* at time *t* and *d*_0_ denotes the background death rate.

### Population Dynamics – P_*i*_

To account for logistic growth of individual populations, our simulation evaluates the impact of density and carrying capacity (P_*i*_) within the dynamic death rate calculation (equation 1). Here, P_*i*_ is determined through species-specific (*S*) population sizes and carrying capacities thus ensuring P_*i*_ = P_*S*_ for all individuals *i* belonging to the same species *S*:

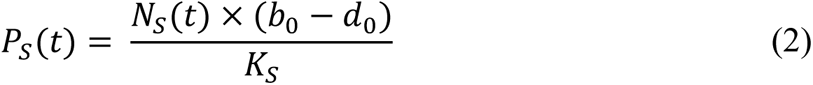

Where *d*_0_ and *b*_0_ represents the background rate of death and birth respectively. *K*_*S*_ and *N*_*S*_ (*t*) represent the carrying capacity and population size at time *t* for an individual of species *S*. The higher the value of P_*S*_, the higher the dynamic death rate of individuals of species *S*.

### Impact of Environment and Trait -*E*_*i*_

The maladaptation of an individual with trait μ at location (*x*, *y*), *E*_μ_ (*x*, *y*) is related to the difference between the individual’s trait value μ and the optimal trait value, or the environment at its location (*u*_0_(*x*, *y*)):

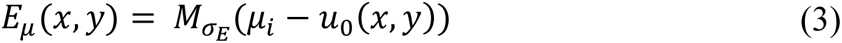

The larger *E*_μ_(*x*, *y*), the worse the individual is adapted to its local abiotic conditions and the higher its dynamic death rate will be. *M*_σ*E*_(*Z*) is a Gaussian function that determines the effect of being maladapted in the current environment. This maladaptedness is weighted by the model parameter σ_*E*_ which controls the severity by which the difference between an individual’s trait and their environment affects the dynamic death rate:

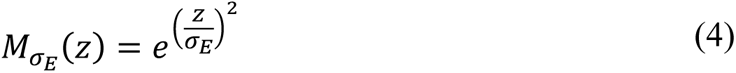

The optimal phenotype, or *u*_0_(*x*, *y*), is determined via a simulation-user defined function. For the study presented here, we assigned a linear function of the one-dimensional environmental gradient (see Figure S1 for a visual representation) but note that this framework can be adapted to more complex landscapes by simply changing the following formula:

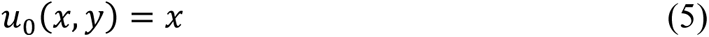

By evaluating environmental maladaptedness as an outcome only of the *x* dimension our simulations enable the formation of alternative community assemblies where species of similar environmental preference, but with competitive association between them may both persevere.

### Species Interactions/Associations –Ω_*i*_

In our framework, the impact of species interactions/associations an individual experiences is a product of the number and species identity of individuals within an area around the target individual. These interactions vary between species according to network matrices.

Our framework quantifies the effect of interactions Ω_*i*_ for a user-defined distance around each individual *i* as follows. The interaction coefficient, *l*_*i*,*j*_ denotes the impact of species *j* on species *i* drawn from the *i X j* network matrix. To account for effects of neighbouring individuals by their abundance, the interaction coefficients are multiplied by the respective abundances of the *j*-th species in the user-defined area surrounding the *i*-th individual (*a*_*j*_). The resulting sum of abundance-weighted interaction coefficients is subsequently standardised by dividing its value by the number of individuals in the *i*-neighbourhood (*a*_*j*_):

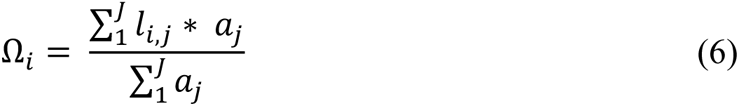

Positive values of Ω_*i*_ reflect beneficial interaction effects and thus decrease the dynamic death rate while negative Ω_*i*_ values increase the dynamic death rate.

### STOCHASTIC SIMULATION ALGORITHM

Our simulation framework tracks birth and death events of individuals belonging to interacting species through time and space as directed by Gillespie, 1977. At each time step *t*, the simulation framework computes the dynamic death rate *d*_*i*_(*t*) for each individual as described above. A birth or death event is selected to occur at random (demographic stochasticity), weighted by the individual’s static birth and dynamic death rates.

If a death event is selected, the chosen individual is removed from the population. When a birth event occurs, the offspring is a clone of the parent individual in their trait value μ_*i*_. Thus, reproduction in our framework is asexual. Natal dispersal is the only form of movement in our framework with offspring location coordinates (*x*, *y*) given as the parental *x* and *y* location, with the addition of a dispersal magnitude drawn from a normal distribution with mean equal to 0 and a user-defined standard deviation. The boundary conditions of the continuous landscape in either dimension are defined by the simulation user. Any dispersal attempt beyond these limits will result in dispersal to the limit.

Next, the time until the subsequent event is determined by randomly sampling from an exponential distribution with a mean of the inverse of the sum of the probabilities all possible events. This is done to ensure that simulation time represents realistic passage of time in biological systems (i.e., more individual birth and/or death events happen in shorter time in larger populations). Finally, the simulation starts over with the updated pool of individuals. This framework is similarly developed in Vinton & Vasseur, 2020.

Lastly, simulations are initialised with a user-defined or randomly generated (1) network matrix, (2) table of initial individuals containing spatial location and trait values for each individual, (3) a vector of carrying capacities for each species, and (4) a function which defines how coordinates relate to environmental values/optimal phenotype (like in equation 5).

### IMPLEMENTATION IN THE R ENVIRONMENT

We coded and executed our simulation framework in R version 4.0.2. (R Core Team, 2021). The simulation inputs (see below) parameterize the simulation framework described above. To ensure reproducibility, we set a seed for random computation processes. For faster computation of several simulation instances at once we enable parallel processing of several simulation runs using the doParallel (Microsoft & Weston, 2020a) and foreach (Microsoft & Weston, 2020b) libraries in R and carry out updates to dynamic death rate values only to those individuals who have been birthed or who have experience a birth or death event in their *a*_*j*_-neighbourhood.

### Network Matrices

Using the randcorr R package (Schmidt & Makalic, 2018), we created supporting functionality with which we establish a set of 1000 random association matrices containing 30 species with a maximum strength of association of 10 to create networks in which associations dominate birth and death probabilities over environmental effects and a matrix-sparsity of 10% to simulate well-balanced networks of diverse association motifs. We focus here on association matrices as most network inference approaches to date aim to infer undirected links.

### Initial Individuals & Carrying Capacities

Using each random network we have created, additional functionality of our simulation framework creates a user-defined number of individuals (either in total or for each species) and allocates trait values to each individual which are drawn from a random distribution around a randomly selected trait mean for each species given a user-defined standard deviation for these random trait distributions. Additionally, individuals are placed randomly within a user-defined spatial extent in x and y dimensions. Lastly, carrying capacities for each species are chosen from a uniform distribution with user-defined boundaries.

For the analyses presented here, we chose random trait mean values between 0 and 10 for each species and set carrying capacities of 300 for all species. We subsequently initialised each simulation with 500 individuals per species in random locations ranging between 0 and 10 in both x and y dimensions. Individual trait values were drawn from random distributions around individual-species-specific trait mean values and a standard deviation of one.

### Simulation Parameters

To use the initialising data (i.e., network matrices, data tables of individuals and carrying capacities) to generate spatially explicit products ready for network inference, we executed our simulation framework with the following parameters. Background rates of birth (*b*_0_) and death (*d*_0_) were set to 0.4 and 0.6 respectively. To calculate local environmental effects on the dynamic death rate, we defined σ_*E*_ as 2.5 (a value at which we found resulting species distributions to be spread, on average, enough for substantial overlap so that species associations could be realised between species-pairs) and supplied a linear function which translates x-values directly into environmental values/optimal phenotypes (see equation 5). When calculating association effects on the dynamic death rate, we defined the size of bounding box around each individual by which association effects and respective species abundances are calculated as 0.5 resulting in a quadratic extent where each side measures 1 unit in x and y dimensions. This was done to correspond with later gridding of the continuous space for network inference although not aligning strictly with each other. Dispersal distance of individuals originating from a birth event were randomly drawn for each dimension (*x* and *y*), for each birth event, from a normal distribution centred on 0 with a standard deviation of 0.2. Finally, we ran each simulation until simulation time 20 was reached, recording individual locations and trait values in time intervals of 0.1 (see Figure S2 for a visual representation of how these individual-time-step information can be used for studying time-series of abundance which a number of network inference approaches may use; e.g.; Damgaard & Weiner, 2021).

### INFERENCE APPROACHES

We carried out ecological network inference using two approaches – HMSC and COOCCUR. We chose these two approaches for their unique placements along a co-occurrence to performance spectrum which characterises network inference input data types and which has been demonstrated to affect network inference outcomes (Kusch et al., 2023). While COOCCUR incorporates co-occurrence data, HMSC can consider species performance implicitly through abundance products as well as explicitly through, for example, population growth rates. Additionally, while COOCCUR is blind to environmental gradients and preferences, HMSC can identify these as part of its inference approach. Lastly, COOCCUR has also been part of a previous simulation-based assessment (Lavender et al., 2019).

### COOCCUR

COOCCUR is a probabilistic framework first introduced by Veech, 2013 which we implemented in our analyses via the cooccur R package (Griffith et al., 2016). COOCCUR identifies species associations in sign and strength through comparison of analytical estimates of species co-occurrence probabilities to purely random species distributions and resulting co-occurrence patterns. Probabilities of observed co-occurrence rates being higher or lower than expected at random are used to denote statistical significance of identified associations. Using these p-values and the traditional cut-off of 0.05, we retained only those associations inferred with statistical significance, setting all other associations in inferred networks with COOCCUR as absent associations for the sake of inference performance assessments.

### HMSC

We executed the HMSC method (Ovaskainen et al., 2017) through the hmsc R package (Tikhonov et al., 2021). HMSC is designed as a hierarchical species distribution modelling framework which aims to explain inputs of site-by-species matrices through environmental niches of species and site-specific environmental characteristics, functional traits of species, and phylogenetic relatedness. Residual covariance of species is then treated as association inference (Ovaskainen et al., 2017). The statistical significance of these inferred associations is available via the credible intervals of their posterior distributions (Opedal & Hegland, 2020). For all subsequent analyses, we only retained those associations per HMSC model which were identified with 95% credible interval not overlapping 0, again assigning absence of association to all remaining inferred associations. To study the effect of informing network inference with environmental conditions, we executed HMSC models in two modes – naïve and informed which represent no consideration of environmental information and consideration thereof, respectively. Furthermore, to assess the effect of data inputs along the co-occurrence-performance spectrum, we executed each naïve and informed HMSC models with three different sets of spatial inputs corresponding to occurrence, abundance and performance data.

### DATA PREPARATION FOR NETWORK INFERENCE

To prepare the simulation outputs for network inference, for each final spatial output, we extracted trait means for each species which correspond to species-but not individual-specific optimal location along the *x* dimension. We then identified species abundances across the continuous two-dimensional landscape in 100 evenly spaced grids (i.e., 10 segments along each axis resulting in grid cells with a length of 1 on either side). Correspondingly, we also extracted for each grid cell population growth (an indicator of species performance) as the change in abundance between the last two simulation time steps. This information was subsequently stored as a species-by-site abundance matrix (see Figure S3 for a visual representation) as well as a species-by-site performance matrix. Lastly, we noted the coordinates of each grid cell and assigned a unique ID to each grid cell. This product was used as a study design identifier and to establish random effect levels with respect to cell ID in HMSC models. For COOCCUR as well as occurrence-informed HMSC analyses, the species-by-site abundance matrix was transformed into a matrix of presence/absence.

### INTERACTION/ASSOCIATION REALISATION

As a product of the random generation of environmental preferences and association motifs per species, some of the simulated associations may never be realised due to association partners not being able to survive in each other’s presence due to their environmental preferences. As such, the simulated ecological networks represent “potential” association patterns which network inference approaches cannot reasonably be expected to identify. Therefore, we distinguish between true potential ecological networks (i.e., the randomly generated input networks for our population dynamics simulations) and true realisable ecological networks. The latter are established by assigning as absent (i.e., magnitude of 0) all associations whose association partners exhibit environmental niche preferences (i.e., initialising mean per species in data simulation) further apart than the radius of the association effect bounding box (i.e., 0.5) plus the niche breadth parameter (σ_*E*_; i.e., 2.5). At a carrying capacity of 300 and a species abundance of 1 without any associations present, an individual 3 units removed from its optimal environment receives a dynamic death rate of ∼1 (∼1.6 at species abundance 300) as compared to the static birth rate of 0.4, thus making death of this individual much more likely than survival. Thus, this condition ought to adequately mask associations which could never affect their involved species-pairs. See Figure 2 for a visual representation of how this process works.

### PERFORMANCE ASSESSMENT

To facilitate quantification of network inference accuracy, we identified whether individual true realisable associations were inferred correctly. Since species may go extinct throughout our simulation runs either due to maladaptation to the environment or competitive exclusion in their preferred environment, we focus on the dissimilarity in links between nodes shared by the true realisable networks including only extant species by the end of the simulation procedure and their inferred counterparts. Expressing the rate of correctly inferred associations divided by the total number of possible links between nodes of the final species pool allows us to express interaction accuracy as a proportion thus making it comparable across different simulation settings and inference approaches. We multiply these scores by 100 to express them as intuitive percentages of whole-network inference accuracy.

To assess sub-network inference accuracy with respect to association identity (present/absent) and sign (negative/positive), we also calculate rates of (1) correct, (2) incorrect, and (3) failure of identification of positive, negative and absent associations.

Finally, we provide a generalisable framework with which to quantify inference behaviour of any network inference approach following our simulation procedures. To do so, we establish two sets of logistic models using the brms R package (Bürkner, 2021) which model two target variables as an outcome of the difference in trait expression per pair of association partners as well as the true strength of associations. These two target variables are each split according to association types (i.e., negative, positive, absent) and denote (1) inference and (2) detection (i.e., correct inference) of individual associations. In doing so, we generate valuable insight into what conditions lead to individual network inference approaches inferring a specific interaction type and under which conditions such inference can be assumed to be accurate.

## RESULTS

Our simulations yielded 1000 distinct random association networks containing 30 species each with random mean trait/phenotype expressions per species per simulation product. These are accompanied by their corresponding 1000 lists of data tables containing individual IDs, phenotypes/traits and locational information which we use for network inference. Here, we focus first on a reduced conceptual simulation run to demonstrate the functionality of the data generation simulation framework and subsequent analysis procedures before we explore validation efforts of network inference at scale.

## SIMULATION OUTPUTS

To demonstrate our simulation framework, we established a conceptual random association matrix containing five species (with a maximum association strength of 10 and a network sparsity of 0.1). See Figure 2 for a visual representation of the conceptual true potential association network matrix as well as the process by which we establish an understanding of the true realisable version of this network. The true potential random association network matrix is then used to calculate the interaction effect on dynamic death rates at each simulation step and shapes the final spatial product of species abundances across the artificial continuous landscape as can be seen in Figure 1.

**FIGURE 1.**
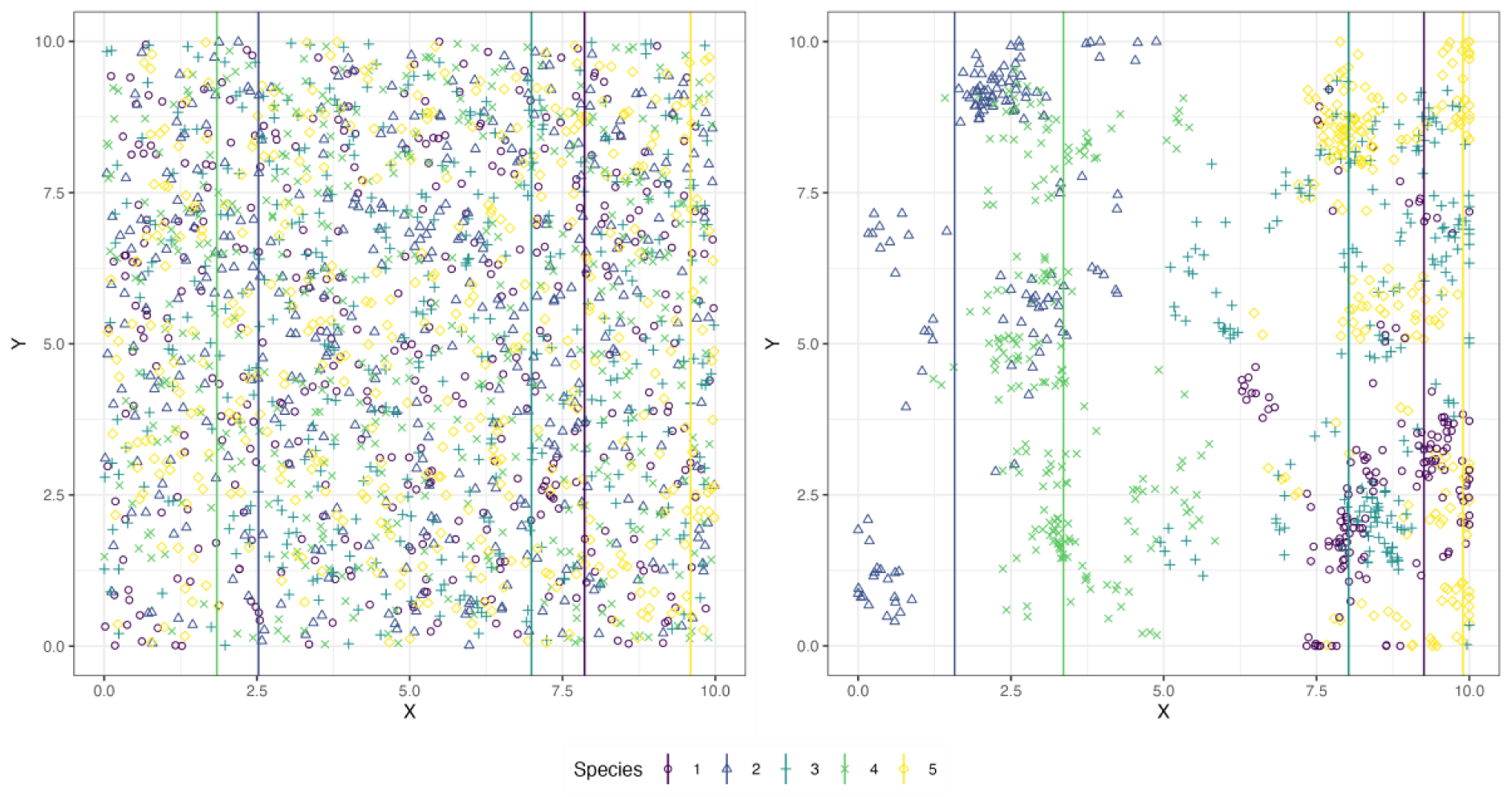
– CONCEPTUAL SPATIAL PRODUCTS. Simulations are initialised (left panel) with individuals belonging to interacting species which are placed randomly across the continuous landscape. This placement is irrespective of trait expressions specific to each species (coloured lines showing the mean trait expression per species in either panel). The simulation terminates at a user-defined simulation-time and returns the final output of individuals across the landscape (right panel) which have shifted with respect to (1) population dynamics, (2) individual environmental maladaptation, and (3) interaction effects.

**FIGURE 2.**
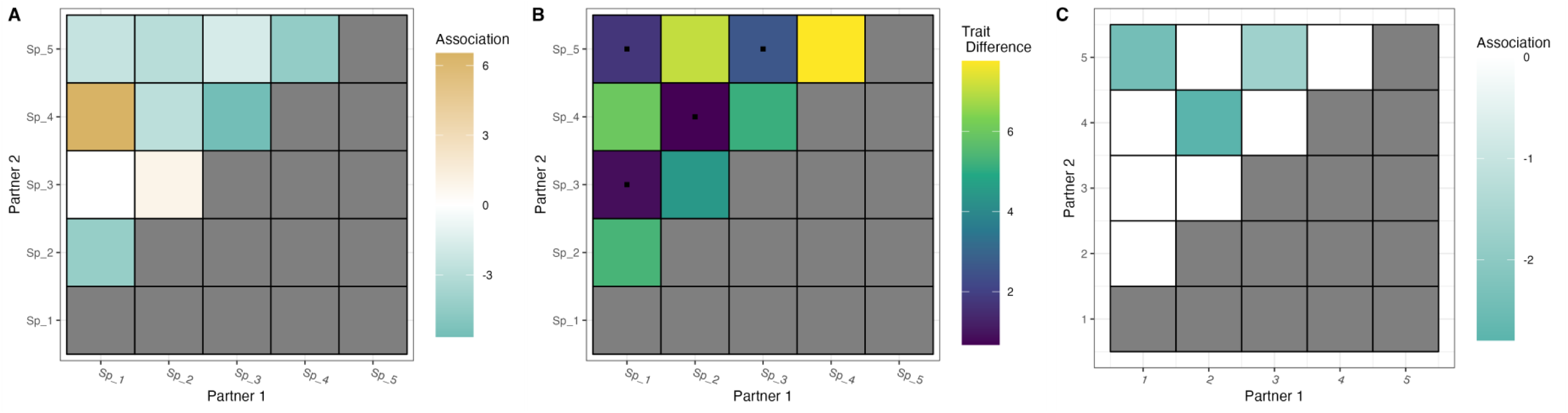
– CONCEPTUAL RANDOM NETWORK MATRIX AND REALISATION OF ASSOCIATIONS. A) A true potential association matrix for five distinct species. Note that maximum association strength was set to 10 and network sparsity to 0.1 resulting in 1 out of the possible ten associations having a strength of exactly 0 while all other associations may range between –10 and +10. B) The pairwise difference in environmental difference as identified through mean and standard deviation of species-specific trait values at the final simulation step (see vertical lines in left-hand panel of Figure 1). Black squares identify associations which can be realised due to shared environmental preferences. C) The true realisable association matrix corresponding to A). All associations which cannot be realised have been set to a magnitude of 0 (i.e., absent associations).

If desired, our simulation framework also returns products containing all extant individuals as well as their traits and locational information at pre-defined simulation-time intervals. This enables the analyses of time-series of species abundances as shown in Figure S2.

### PERFORMANCE OF NETWORK INFERENCE METHODS

Using COOCCUR and HMSC in our conceptual example, the inferred networks are notably different to the true, original networks which we used to drive our simulations (Figure 3) with the exception of absent associations in most HMSC inferences.

**FIGURE 3.**
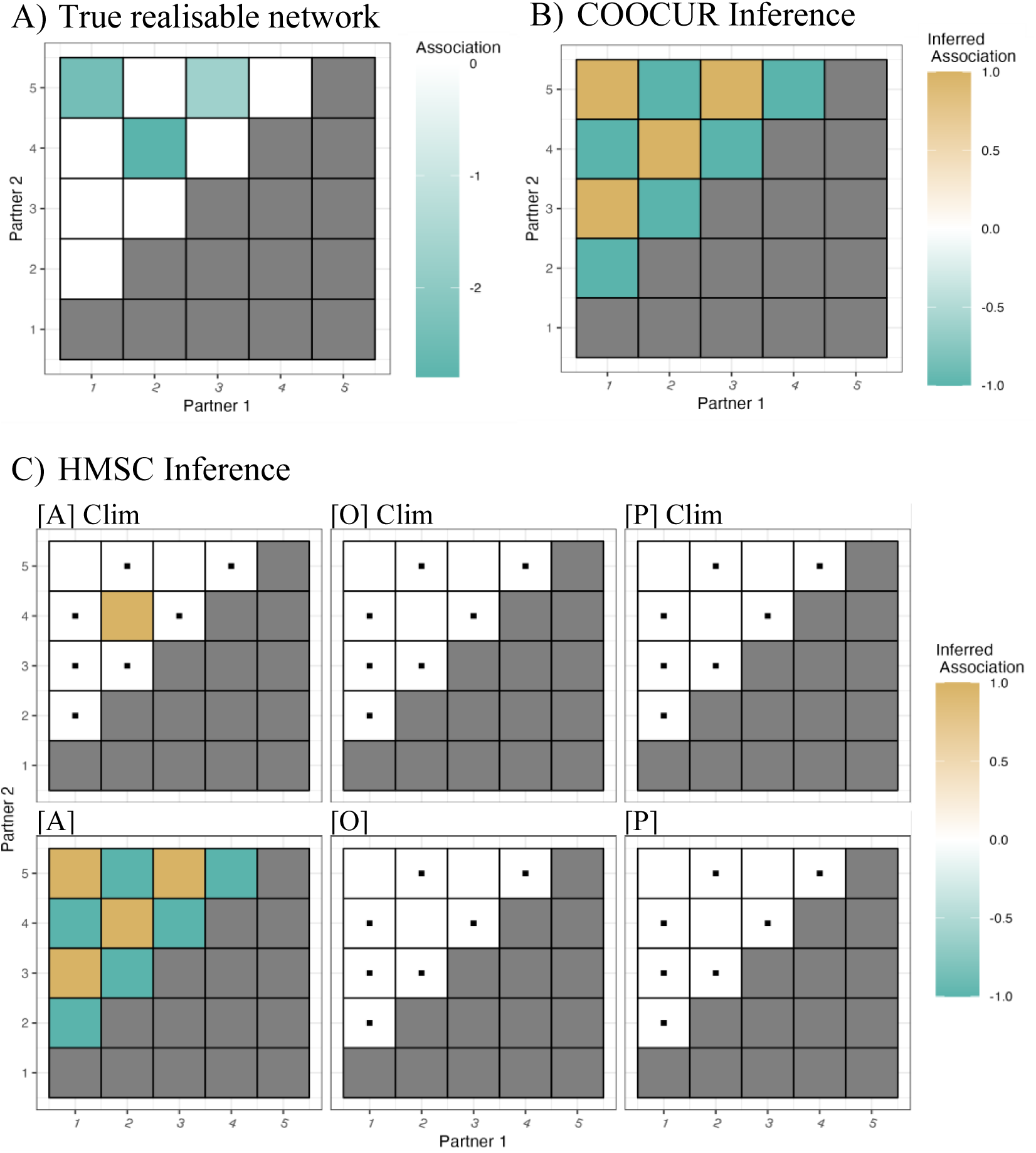
– CONCEPTUAL INFERRED NETWORKS. Inferred networks of statistically significant pairwise associations (whose association strengths have been set to –1 and +1 respective to association inference sign) show great dissimilarity to original network (figure 2, c) which affected the simulation framework. Black squares identify correctly identified positive, negative, and absent associations. Abbreviations: O; A; P denote co-occurrence, co-abundance, and co-performance data inputs. Clim identifies models informed with environmental data. Black squares denote correctly inferred associations.

Assessing network inference accuracy for all 1000 simulation outputs of 30-species networks, we find that, within our analyses network inference accuracy varies largely across our two network inference approaches and their sub-specifications (Figure 4). We find that COOCCUR performs worse (a mean average network inference accuracy of roughly 23%) and is outdone by HMSC inference irrespective of how HMSC models are specified or informed. However, within HMSC, we find that network inference accuracy is governed by spatial input type and environmental parameterisation.

**FIGURE 4.**
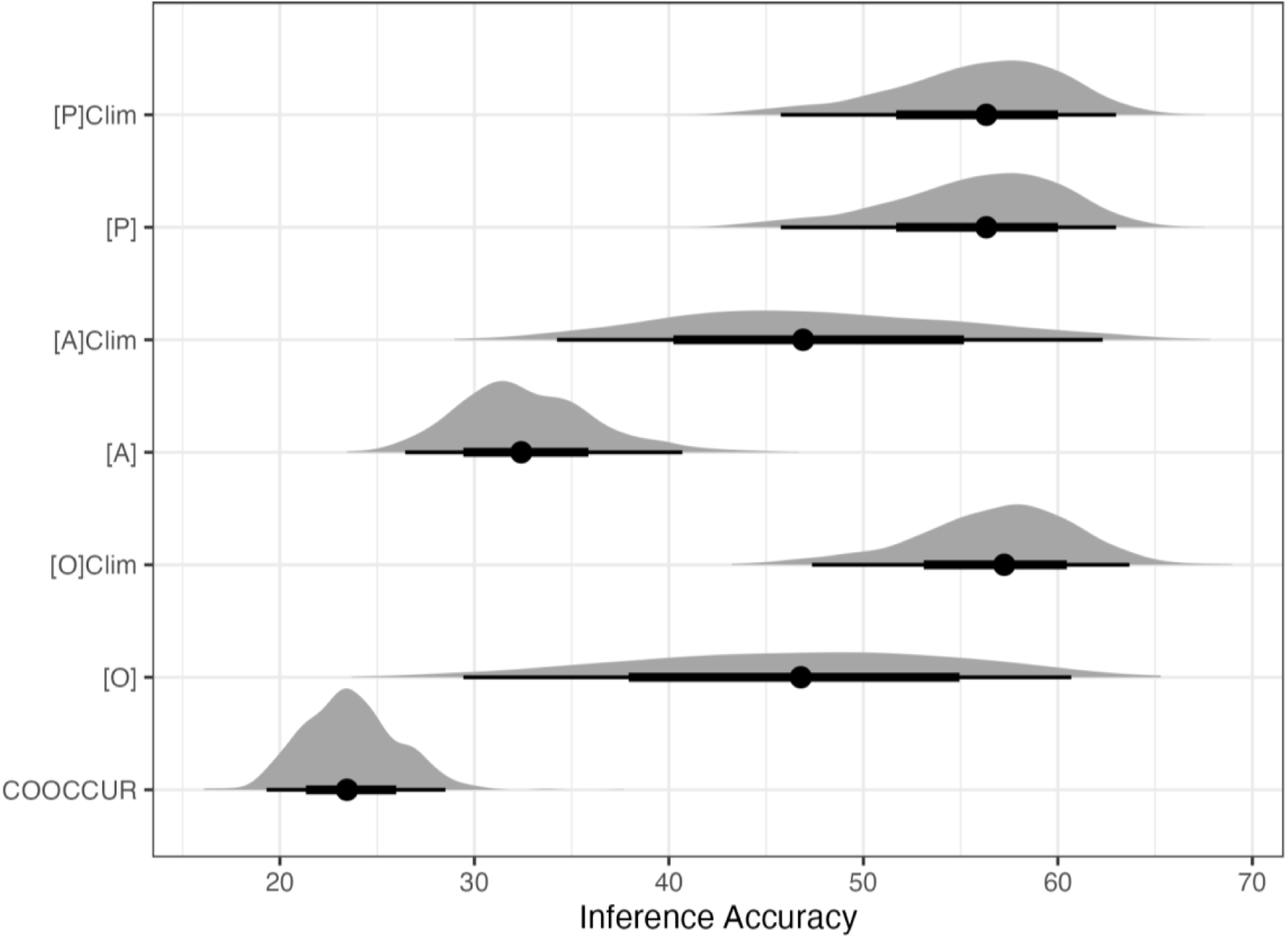
– NETWORK INFERENCE ACCURACY. When analysing similarity links of shared species between inferred networks and the true realisable association networks for each inference approach across all simulation runs, we find that COOCCUR produces less accurate inference than HMSC. Meanwhile, HMSC specification and inputs determine HMSC network inference accuracy. Abbreviations: O; A; P denote co-occurrence, co-abundance, and co-performance data inputs. Clim identifies models informed with environmental data.

While we find that inclusion of environmental information in HMSC models generally leads to greater network inference accuracy, this effect is not present when utilising co-performance inputs. Similarly, while we find a general trend of greater network inference accuracy for HMSC models when moving from occurrence to abundance and performance data, HMSC models informed with abundance data but not accounting for environmental effects perform substantially worse than other HMSC specifications. Most strikingly, however, we find that occurrence-based HMSC models which account for environmental effects perform on-par with performance-informed HMSC models. For a breakdown of network inference similarity across approaches and HMSC specifications see Figure S4.

Further sub-network deconstruction of network dissimilarities into rates of true, false and not-inferred (i.e., missed) positive, negative and absent associations identified in each inferred network (see Figure S5) under our simulation settings reveals that (1) COOCCUR and HMSC (irrespective of specification) perform comparably at correctly identifying positive and negative associations with both approaches struggling to correctly identify negative association, (2) HMSC, irrespective of specification, performs better at correctly identifying association absence than COOCCUR, (3) both COCCUR and HMSC infer positive, negative and absent associations incorrectly at high rates with absent associations assigned by COOCCUR being wrong at markedly higher rates than those rendered by HMSC inference, (4) COOCCUR and abundance-informed HMSC models fail to identify positive and negative associations the least. Meanwhile, (5) these two specifications fail at inferring true, realised absent associations and are far outdone by HMSC model informed through performance data. This highlights a stark contrast in network inference reliability given the underlying true realisable association type.

## NETWORK INFERENCE APPROACH BEHAVIOUR

To further understand network inference approach behaviour, our logistic regression models using the difference in trait expression per pair of association partners as well as the strength and sign of associations to predict the probability of (1) assigning (i.e., inference) and (2) assigning correctly (i.e., detection) either positive, negative, or absent associations to species-pairs for COOCCUR and HMSC. For clarity and brevity, we focus only on models of positive association inference and detection using COOCCUR as well as abundance-informed HMSC models due to their clear distinction in average network inference accuracy (Figure 4).

## INFERENCE PROBABILITY OF ASSOCIATIONS

We find that the inference of positive associations between species pairs is unaffected by the true realisable association between species pairs for COOCCUR as well as abundance-informed HMSC models. However, we do find that decreasing difference in niche preference of species-pairs (which corresponds to their separation in space) leads to a rapidly increasing probability of inference of positive associations (Figure 5). While this pattern persists across all three network inference approaches assessed here, this behaviour is strongest for COOCCUR, followed closely by the naïve implementation of abundance-informed HMSC while the informed HMSC model shows the weakest signal of niche-preference differences.

**FIGURE 5.**
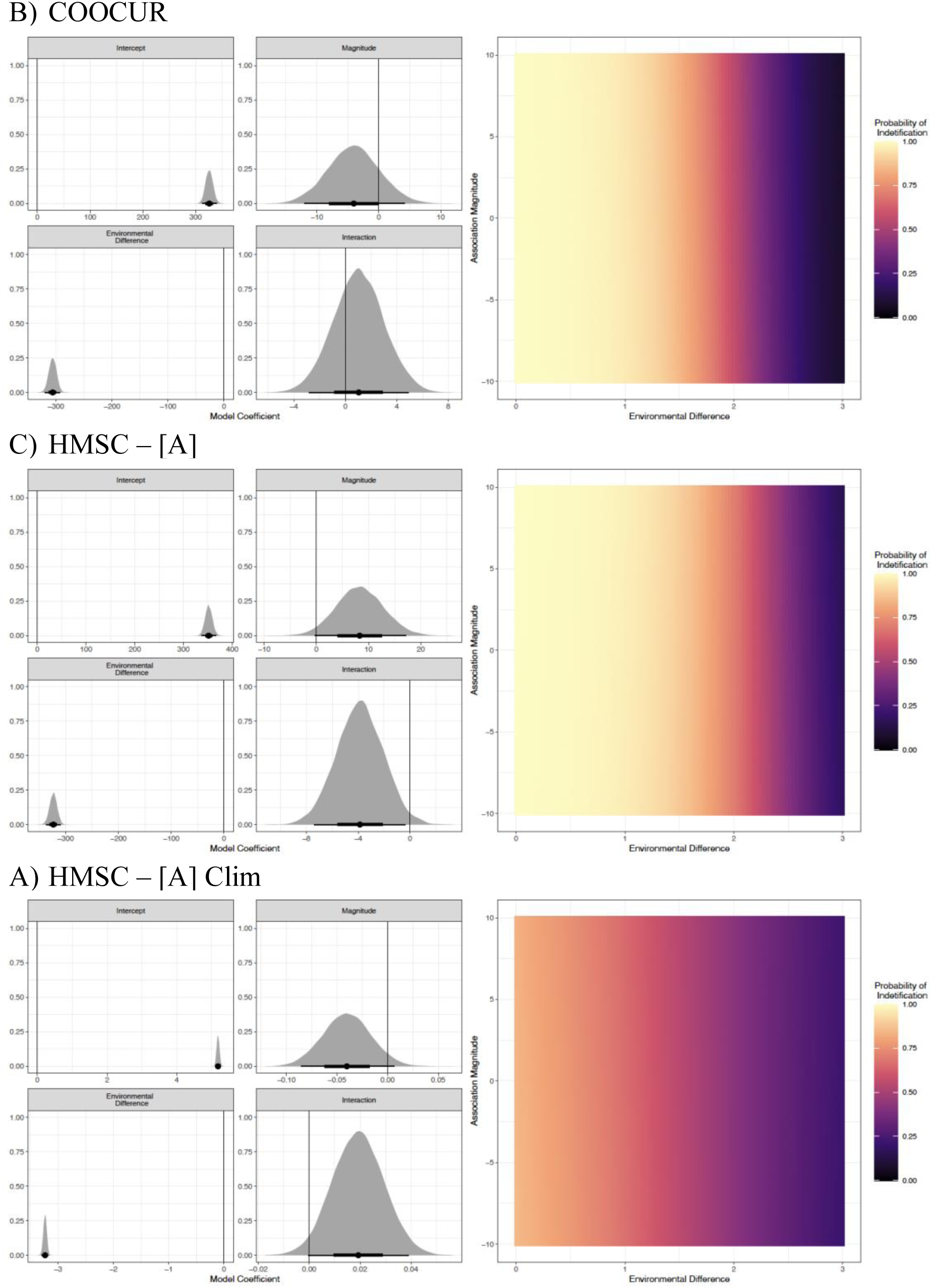
– PROBABILITY OF CORRECT IDENTIFICATION OF TRUE REALISED ASSOCIATIONS. Logistic regression coefficients (left column) show clear effects of environmental difference in species which translate to distinct probabilities of inferring a pairwise associations (right column). However, this effect is diminished for abundance-informed, environmentally-informed HMSC models as compared to their naïve HMSC counterparts and COOCCUR. Abbreviations: A denotes co-abundance data inputs; Clim identifies models informed with environmental data.

## DETECTION PROBABILITY OF ASSOCIATIONS

Detection of positive associations is driven, for the three evaluated network inference approaches, by a combination of the magnitude of the underlying true realisable positive association as well as environmental niche preference difference of species-pairs. Detection probability increases with underlying association magnitude and decreasing environmental niche preference differences (Figure 6). All three approaches behave similarly as was to be expected from their indistinguishable true positive association detection rates (Figure S5).

**FIGURE 6.**
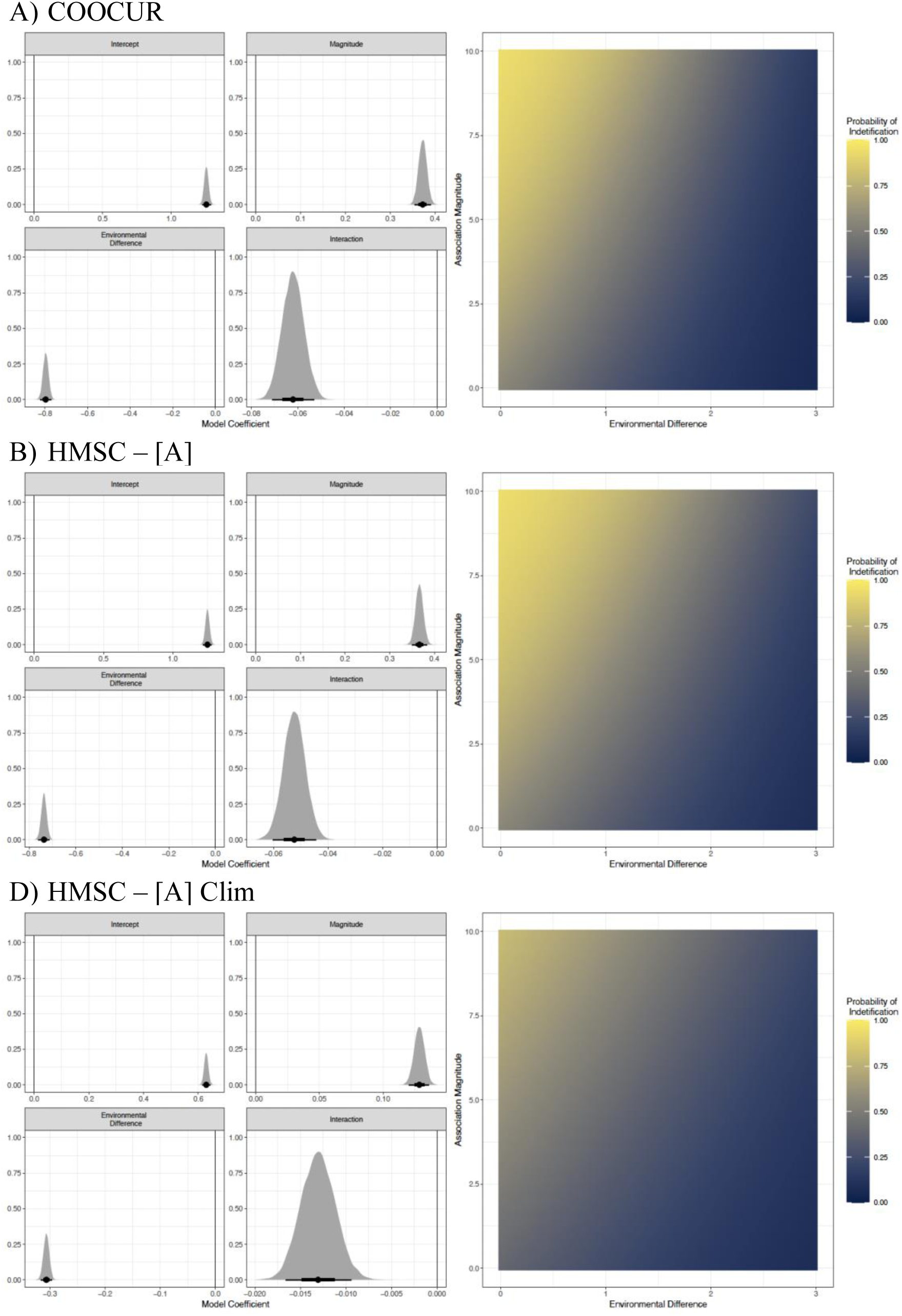
– PROBABILITY OF CORRECT IDENTIFICATION OF TRUE REALISED ASSOCIATIONS. Logistic regression coefficients (left column) show clear effects of association magnitude, environmental difference in species niches as well as the interaction of these two which translate to distinct probabilities of correctly identifying a pairwise associations (right column). Abbreviations: A denotes co-abundance data inputs; Clim identifies models informed with environmental data.

## DISCUSSION

Aligning with previous recommendations for choice of network inference approach (Kusch et al., 2023), we find that network inference accuracy is governed by (1) spatial biodiversity input type along a spectrum of co-occurrence, abundance, and performance which confers increasing accuracy onto networks inferred therefrom, and (2) environmental considerations in network inference approaches with network inference approaches which account for environmental conditions rendering more accurate inferences of true networks than those approaches who don’t. These findings confirm previous criticisms of co-occurrence based network inference approaches failing to capitalise on existing information in abundance and performance data (Bimler et al., 2022; Kusch et al., 2023; Ovaskainen et al., 2017). In addition, our findings also highlight the previously suggested fallacy of neglecting environmental gradients in network inference leading to erroneous assignment of positive and negative because of differences in environmental preferences rather than actual interactions (Blanchet et al., 2020). This is particularly demonstrated via our inference model outputs in Figure 5 where inclusion of environmental information leads to a diminished impact of spatial separation on inference likelihood of positive associations. Lastly, we also find that the underlying inference approaches themselves (i.e., COOCCUR and HMSC) affect accuracy of network inference with both naïve occurrence-informed HMSC models and COOCCUR using the same inputs, but HMSC performing far better at accurately inferring ecological networks (Figure 4). However, these findings do not hold for all approach specifications in our simulation set-up as described here. Notably, there seem to exist saturation effects to inference accuracy gains with there being no difference in inference accuracy of naïve and informed performance-HMSC models as well as informed occurrence-HMSC models. While this may suggest that the inclusion of either performance data or environmental data may suffice in maximising network inference accuracy, the markedly worse performance of naïve abundance-HMSC models when compared to their informed abundance-HMSC counterparts should serve as a cautionary tale to neglecting environmental data inclusion.

While assessing whole-network inference accuracy revealed a clear and strong effect of choice of network inference approach as well as input data selection, less generality can be identified at sub-network scale. We find that different inference approaches and specifications thereof have different likelihood of rendering correctly positive, negative, and absent associations. Absent associations are particularly well-rendered by performance-HMSC models. However, the assignment of absent associations for these models was so frequent that they failed largely to identify associations belonging to other types. Meanwhile, all other specifications of HMSC as well as COOCCUR perform similarly well at detecting true positive associations, but also infer their presence incorrectly at high rates (Figure S4). Informed abundance-HMSC models performed the best at detecting true negative associations, but do so nevertheless at worryingly low rates of success. Owing to this varied association-type-specific detection performance of our tested approaches, we suggest that it may be useful to establish inference of ecological networks via ensemble approaches which prioritise association types inferred with one approach over the same association type inferred by different approaches. Caution should be exerted in transferring these findings for use in other studies however as our simulation settings will affect these performances. Thus, for judging how to best build an association-type specific network inference ensemble approach, it is pivotal that network inference performance is first validated with simulation parameters approximating study-specific conditions.

We expect that using our simulation framework in conjunction with real-world association networks and bioclimatic niche preferences and environmental suitability constraints will yield a more realistic overview of network inference performances. However, such a study will require extensive data collection to allow more precise parameterisation of the simulation framework we are presenting here. To circumvent the tedious data collection and costly simulation effort this undertaking would entail we propose that the logistic regressions we present may help, in first instance, to investigate under which conditions association detection is likely. Instead of selecting one “best-performing” network inference approach, we believe that it may be more prudent to quantify capabilities of network inference approaches to identify association types with respect to association sign, magnitude, and differences in environmental preferences of association partner-species as we have done with our logistic models in Figure 6. Doing so will create ranges of environmental and biological constellations within which individual network inference approaches can be treated as reliable.

Even so, we do not claim our network inference performance scores nor our logistic regressions to be necessarily representative of real-world applications, but merely that they remain accurate for our specific simulation parameters. For example, increasing σ_*E*_ (i.e., making the environmental constraint more forgiving) or increasing association magnitudes and introducing a lower limit to association magnitude would likely increase the performance of either network inference approach as the effects of associations will become greater in comparison to environmental preference and location-specific environmental mismatches. Additionally, we suspect that increasing the distance which defines the *i* –neighbourhood may make identification of associations more accurate as interaction effects will have greater reach, thus affecting each individual site in the site-by-species matrix used for network inference more than before. Alternatively, restricting interactive effects not to the *i*-neighbourhood but strictly to the grid cells which define sites should improve performance of either network inference approach and be more representative of study setups with distinct sites (López-Segoviano et al., 2021) rather than continuous megaplot designs (Lutz et al., 2012). Such considerations would incur additional development of the simulation framework we present here.

While our simulation framework represents a major advancement of our capability of identifying network inference performance and validate inferred networks, we also realise that the simulation framework we present here may be regarded as an oversimplification of natural processes. For examples, non-natal movement, which is absent from our framework, has been identified as a confounding effect to interactions studies of animals (Blanchet et al., 2020; Matthiopoulos et al., 2015). Subsequently, we expect the current version of our simulation framework to be most useful to studies of plant-plant interactions/associations or those of other sessile organisms. Furthermore, our simulation framework only includes one functional trait per individual despite organisms inhabiting a multi-dimensional space of functional trait expressions (Blonder, 2018). Our framework neglects the impacts of functional trait expression on association/interaction outcomes which have been demonstrated particularly among tree species (Kunstler et al., 2012). Thus, we believe that our simulation may be further enhanced by the inclusion of additional functional trait metrics which determine the dynamical death rate through the interaction/association coefficient (Ω). Additionally, many species reproduce sexually rather than clonally and there may thus exist considerable variation in such functional traits between parents and offspring. Despite these avenues for future development, our simulation framework remains the most realistic for network inference data generation purposes particularly when compared to previous attempts at quantifying pairwise association inference performance (Lavender et al., 2019).

## CONCLUSION

Network inference approaches have been developed to address the labour intensity of ecological network research which requires impractical sampling efforts particularly at large geographical scales or for large species pools. Despite the large number and diversity in approaches of network inference frameworks, no general framework for network inference validation or performance quantification has been established yet. Here, we have presented an easy-to-use, mathematically robust simulation framework which can be used for data generation with which to carry out network inference validation. While we find promising accuracy of network inference under specific conditions (i.e., use of site-species performance matrices and environmental information), we argue that these performance scores may be subject to change given further parametrising of our simulation with more realistic real-world data. Additionally, further explorations of network inference accuracy of directed links (i.e., interactions) or using the capacity of our simulation framework to output time-series of individuals across space may prove particularly useful in aiding to the overview of network inference capabilities and current practices thereof.

Nevertheless, using our simulation framework and a novel network inference performance and behaviour quantification pipeline, we have confirmed previous criticisms of ecological network inference practices and added to existing guidelines for choice of appropriate network inference approaches. Thus, our simulation framework for network inference validation represents an important addition to the toolbox of network inference method development which fills a crucial knowledge gap of network inference performance assessments.

## ACKNOWLEDGMENTS

This research was supported by the Aarhus University Research Foundation Start-up Grant (grant no. AUFF-2018-7-8) to Alejandro Ordonez.

## AUTHORS’ CONTRIBUTIONS

E.K. conceptualised the study. A.V. developed the simulation framework used in this study.

E.K developed the network inference and network validation framework used in this study.

E.K. created all R scripts necessary for analyses. E.K. led the analysis and manuscript writing.

All authors contributed critically to the drafts and gave final approval for publication.

## DATA AVAILABILITY

All code required for data generation, network inference, and inference performance quantification is available at https://github.com/ErikKusch/Scaleable-Inference-of-Ecological-Networks (please note that this is currently a private repository until submission to a relevant journal so that the simulation framework specification remains protected). We are ready to submit code and data to dryad upon acceptance of the manuscript.

