## Supplementary Material for "A Novel Simulation Framework for Validation of Ecological Network Inference"

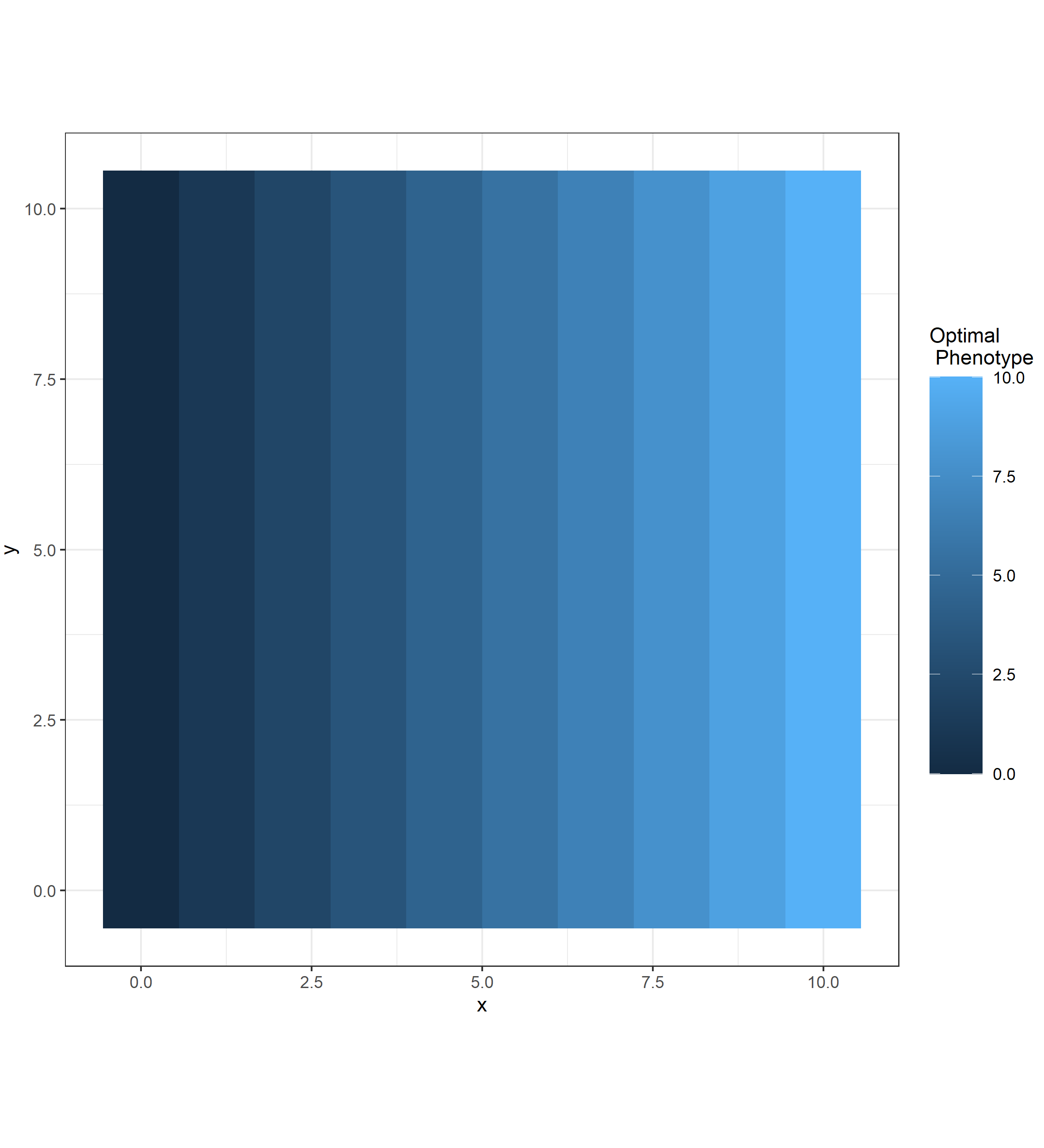

Figure S1 – Spatial Gradient used in simulations. Our analyses were executed using a linear environmental gradient along the x-axis which established the optimal phenotype/trait value for each location of $\boldsymbol{(x,y)}$. Note that this visualisation is exaggerated to highlight how the optimal phenotype is affected only by $\boldsymbol{x}$. The spatial gradient used in the simulation framework is continuous rather than categorical as shown above.

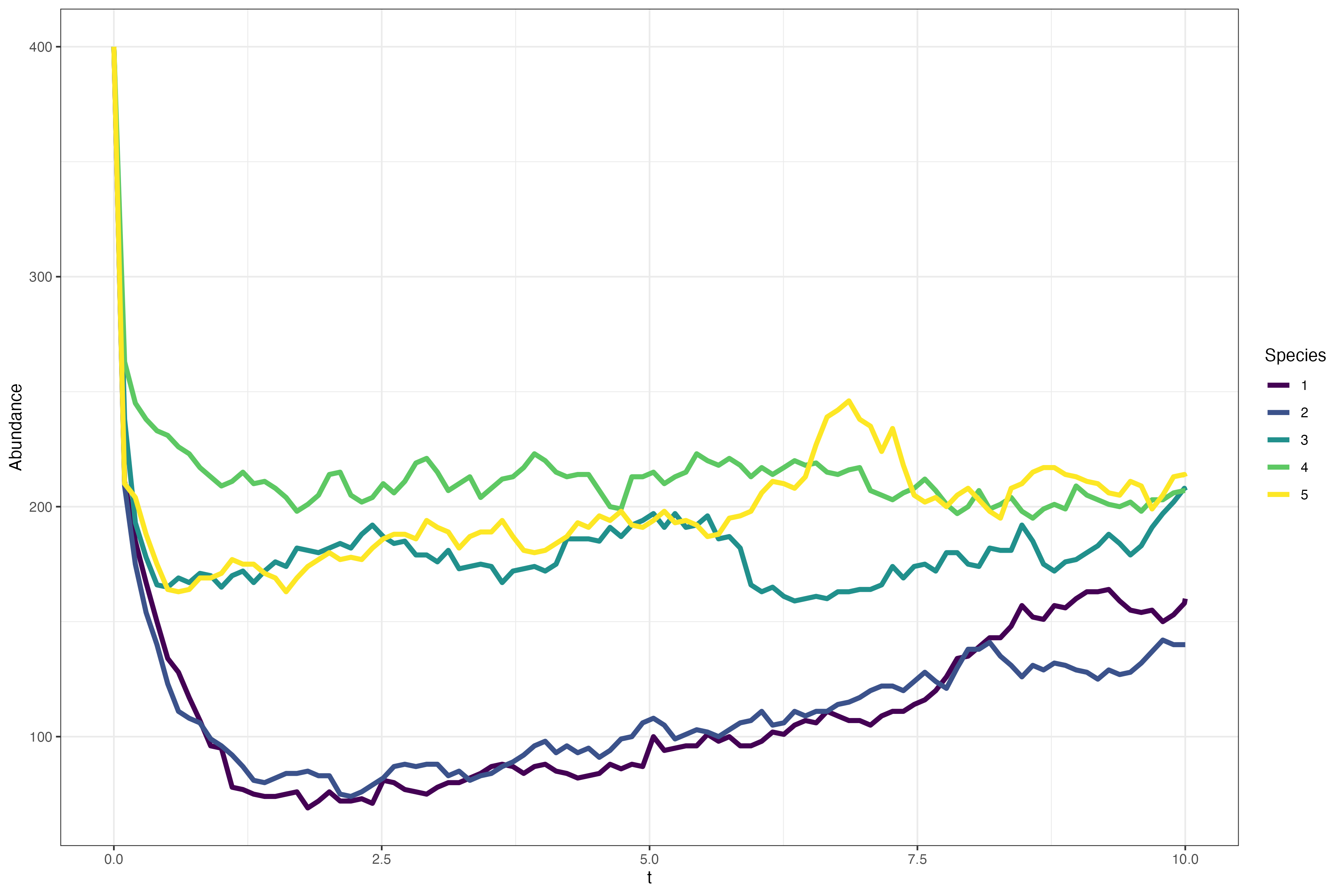

Figure S2 – Abundance of Conceptual Species throughout the simulation run. Our simulation framework can export information on extant individuals throughout the simulation at user-defined intervals with which additional studies can be carried out such as network inference via analyses of time-series of abundances.

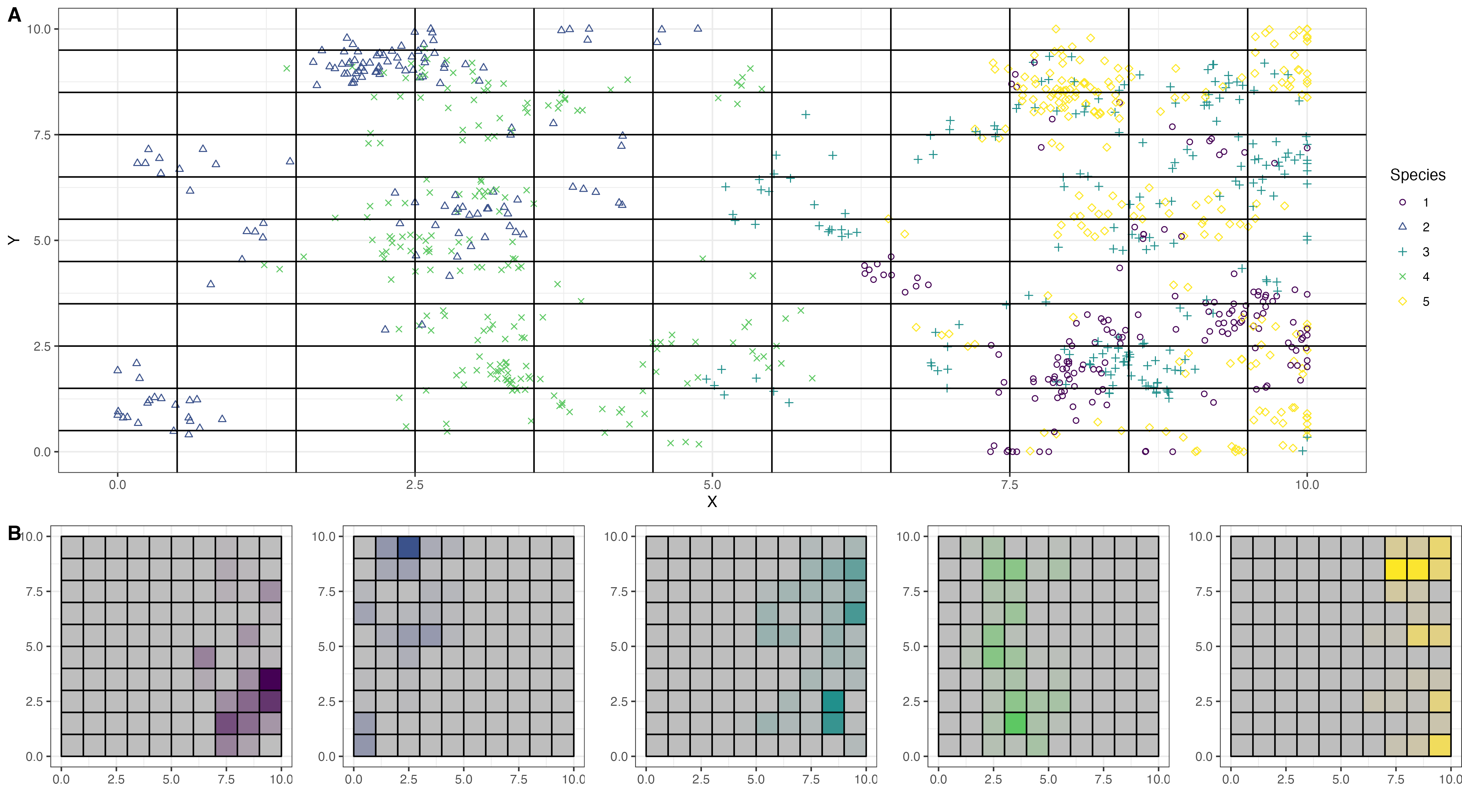

Figure S3 – Conceptual Spatial Product and Matrices for Network Inference. Network inference approaches leverage species-by-site matrices of presence/absence or abundance. To establish these, we divide our continuous landscape (A) into evenly spaced grid cells for which the abundance (B) of each species is then recorded and stored in a species-by-site matrix.

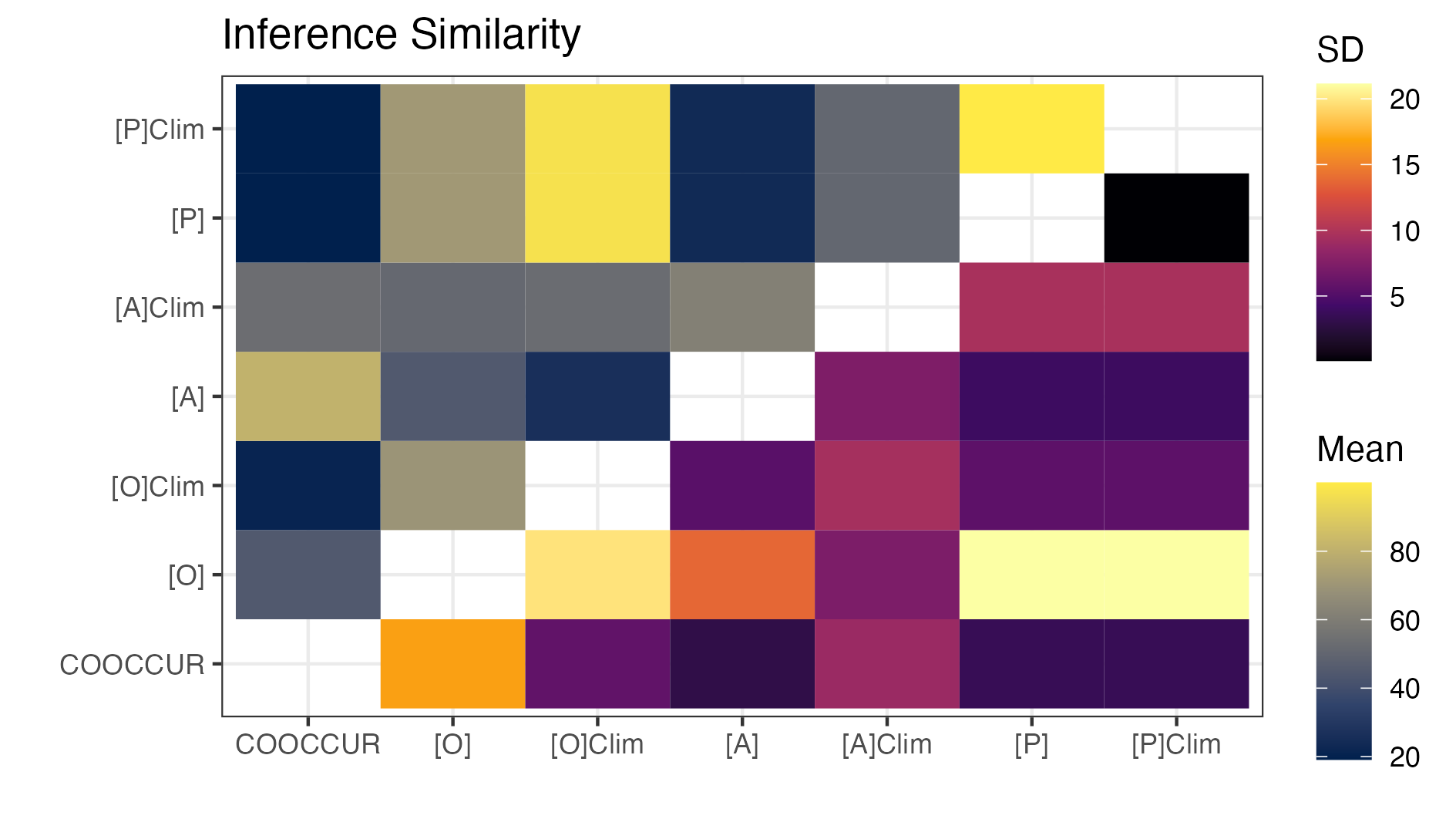

Figure S4 – Dissimilarity of Inferred Networks. Networks inferred by HMSC and COOCCUR per simulation are markedly dissimilar, but strong similarities exist between network inferred using performance-informed HMSC models as well as climate-informed occurrence-HMSC models. Abbreviations: O; A; P denote co-occurrence, co-abundance, and co-performance data inputs. Clim identifies models informed with environmental data.

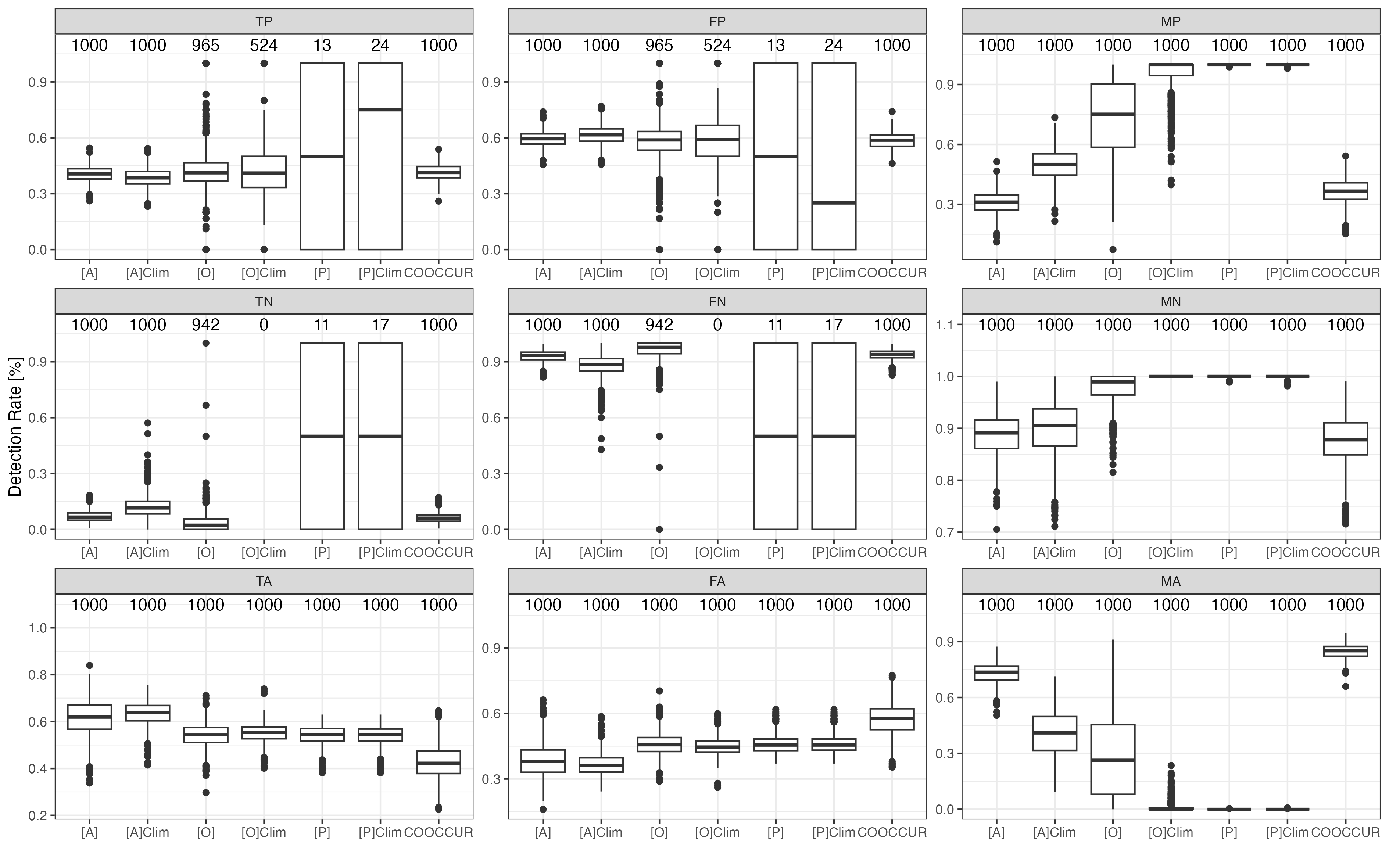

Figure S5 – Rates of True Realised Association Detection. Decomposition of overall network dissimilarities expressed as density distributions reveal similar and low performance of the two network inference approaches for our simulations. Two-letter codes denote type of error rate: TP…True positive, FP…False positive, MP…Missed positive (a positive association that was failed to be identified), TN…True negative, FN… False Negative, MN… Missed negative, TA…True absent, FA…False absent, and MA…Missed absent. Model abbreviations: O; A; P denote co-occurrence, co-abundance, and co-performance data inputs. Clim identifies models informed with environmental data. Numbers denote how many inferred networks could be used for plotting (i.e., how many inferred networks contained at least one of the assessed association inference types).
